# Two biological constants for accurate classification and evolution pattern analysis of Subgen.*strobus* and subgen. *Pinus*

**DOI:** 10.1101/297606

**Authors:** Huabin zou

## Abstract

Currently, biological classification and determination of different categories are all based on empirical knowledge,which is obtained relying on morphological and molecular characters. For these methods they lacks of absolutely quantitative criteria ground on intrinsically scientific principles. In fact, accurate science classification must depend on the correct description of biology evolution rules.

In this article a new theoretical approach was proposed, in which two characteristic constants were gained from biological common heredity and variation information theory equation, when it is at the maximum information states, corresponding to symmetric and asymmetric variation states. They are common composition ratios, *P_g_* =0.61, and *P_g_*=0.70. By analyzing the common composition ratios of compounds among oleoresins, two pine subgenus:Subgen.*Strobus* (Sweet) Held and Subgen. *Pinus* could be integrated into one class, Genus pinus, excellently, when *P_g_* = 0.61.

These two pine subgenus could be classified into two groups clearly,when *P_g_* = 0.70.

The results is somewhat different from that achieved by means of classical classification relying on morphological characters. On the other hand, the evolution relationship of two subgenus was analyzed based on characteristic sequences of samples, it indicated that white pine origin from pinus tabuliformis. The two constants should be used as the classification constants of some biological categories of plants.

## Introduction

Biological taxonomy,or systemics is a very classical research field, and it appears that all taxonomic problems have been perfectly solved so far. However, until recently, there exist no any strictly scientific definition and examination about biological species, genus, family, order, class, phylum and kingdom. Thus, this makes biological science building upon the beach. Currently, there are some species definitions, such as biological species concept (BSC), phylogenetic species concept (PSC), evolutionary species concept (ESC), genetic species concept (GSC)， recognition species concept (RSC)，cohesion species concept (CSC) [1,2,3,4,5,6]. All these species concepts are proposed based on empirical knowledge and subjective inference. In fact the definitions of species and other categories are very fundamental for the research on biological diversity, and are the ultimate goal of systemics. Further more, the scientific definition of species is the crown and the holy grail[7]. In reality, very complex biology phenomenons and behaviors make it hard to put forward a perfect and unified species definition for people. This also leads to the difficulty in biological conservation in practice[8] nowadays.

So far, for the definition of biological species, it lacks support of elemental principle theory based on experimental science and mathematical principles, and lacks theoretically quantitative criteria deduced from these principles. In practice the identification and classification of plants totally depend on a lot of morphological characters[6]. Even in modern numerical taxonomy, the identification standards are also from empirical knowledge, learning/training samples. So, the development of biology is deeply blocked by empirical classification. For these reasons biology is not a rigorous science. In recent classification mathematics, the major methods are some similarity and difference coefficients [9,10], which are short of deterministic criteria for judging species too. There are many pattern recognition methods constructed grounded on statistics, such as systematical cluster, Principal component analysis (PCA)[9,10]. these approaches are able to group organisms into different classes without any prior knowledge related to themselves. But there is no objective or any unsuspected standard for different classes. Some supervised methods, such as Bayesian [6,11] and Fisher determination [9],are certainly able to classify samples, but they must gain their standards relying on learning samples, which only fit to limited sample sets. The maximum likelihood(ML) [4,6,11] classify samples depending on some unproved hypothesis established on experience. It can not make the conclusions undoubtedly. Biological classification science still lacks of precise science principle which likes that existed in physics and chemistry.

It is necessary to search for absolutely quantitative standards, rather than relative criteria and subjective inference standards established by means of empirical knowledge, for classifying different categories and determining accurate evolution relationship among different organisms. Otherwise, one can not discover the true evolution rules of them. The uncertainty in biological classification brings about difficulty to build up a systematically biological science, and influences deep research on biology seriously. The present state of biology is similar to that of chemistry in the early of 19 century, when thermodynamics was not built up, and periodic table of elements was not be discovered, that is, chemistry theory was an empirical system in that time.

The key feature of principle theory is that it can predict matter’s intrinsic properties accurately and quantitatively. However, the strait of theory based on statistical description can only offer probability. Author Zou found that the two characteristic constants derived from biological inheritance and variation information theory equation, that is dual index information theory equation could accurately identify combination Chinese medicines[12,13,14,15], which are very complex biology systems. That is, the two constants are qualified to be the absolutely quantitative criteria of traditional Chinese medicines (TCM), which consist of extracts of several kinds of herbs. So far there is no report whether the two constants are suited to be as the classification criteria of different organisms, and fit to distinguish biological categories, such as species, genus, subgenus, family and so on.

In generally, a certain and undoubted classification system, accurate definition of species, genus and family and so forth are one of the ultimate aims of biological science[7].

In the respect of classifying plants of pinus, the major methods are based on morphological or macro characters[6]. In modern taxonomic field, the classification of pinus’ plants is also grounded on macro characters. That is, the number of vascular bundle within needles. They are divided into two subgenus, haploxylon with unidimensional bundle, and diploxylon with double vascular bundles[16,17]. In 1966, *P. krempfii* Lecomte was discovered, and it was one new subgenus[18], thus pinus includes three subgenus.

Plants of Pinus originated from China are classified into two subgenus, haploxylon and diploxylon[19]also, according to the number of bundles in a needle. Song Zhenqian divided pinus, growing in China, into three subgens too, that is, subgen. *Strobus*(Sweet)Rehd, subgen.*Pinus* and subgen, Parrya (Mayr) Chu et X.P.Li [20].For the third subgenus, it merely contains two kinds of pines.

On the other hand, a series of researches have revealed that the compositions of rosins of different pines can fully reflect their genetic traits,hereditary characters.

According to literature[20], Herty C.H. (1908) once studied *P.elliottii* Englem. and *P. palustris* Mill.. Dupont G and Barrand M (1925) investigated *P.nigra*, Krestinsky V et al (1932) researched *P. sylvestris*. They also found that for the same kinds of pines, the compositions in rosins varied little. Oudin A(1938) studied *P. pinaster* and obtained the same conclusions. For the same pine trees their compositions of rosins varied lightly in different seasons in an year. The study indicated the collecting rosin methods and ages of pine trees did not influence the compositions[21].

Mirov N.T. analyzed the compositions of 77 kinds of rosins corresponding to 77 kinds of pines, and found that they could be used for identifying pines[22]. He also researched the extracts from different kinds of pines.Their extracts differed from each other obviously, and could be used in chemical classification (chemotaxonomy)[23]. In the early of 1960s, chemotaxonomy was put forward[24,25,26], and this method was applied to classifying pines.

Song Zhenqian studied the oleoresins of *P.koraiensis* Sieb.et Zucc (red pines) grew in Dunhua district of Jilin province in China and in the area of Russa in the European, He found that the compositions of oleoresins from the two areas were highly identical, even though these oleoresin samples produced apart from thousands miles away. The compositions of the oleoresins of Zhan pine originated from Fuyuan district of Zhejiang province in China and from the kamaka mountain in Peru were highly similar to each other[20].Thus, the compositions in oleoresins of different pines can provide us with plentifully molecular characters for classifying pines. Generally speaking, compared to genes, substances in metabolites enable to reflect the live physiology states of organisms more perfectly, since genes representation are regulated by many factors. Thus, gene information can not reflect the physiological states lively.

In august of 1988 and 1989, for each kind of pine, 5 to10 oleoresin samples from the 5 to 10 pine trees were collected, randomly from its distribution center by means of the same manner. Totally hundreds of oleoresins samples from 50 trees belong to 24 kinds of pines were studied systematically. He designed a method for Gas chromatography (GC) analysis without advanced separation, all compositions in oleoresins were analyzed successfully.

In terms of classical classification results of pines,Song analyzed oleoresins of 24 kind of pine samples Subgen.*Strobus* and Subgen.*Pinus* originated in China, and performed the chemical classification of these pines. However, there is no Hierachical Cluster Analysis and pattern recognition being carried out by means of mathematical method so far.

For matter, its constructs determine its functions. For this reason the reliable information for classifying samples should be the material structure information, which pose more rigorous independent characters. Compared to morphological characters, molecular construct characters ought to well represent species and evolution traits of biology.

In mathematical classification of biology, whether there exist some strictly certainty principle theory, which can give us certain classification results. In this article grounded on the biological heredity and variation information theory equation proposed by author Zou[12], two characteristic constants, corresponding to symmetric and asymmetric variation states were obtained, when the information values are at the maximum. These constants could be used as the absolute criteria for classifying complex biology systems TCM based on chemical fingerprint spectra[12-15], and the two constants were defined as the species constants of TCM too[15].In fact, the compositions of pine oleoresins can be viewed as chemical fingerprint spectra of compounds. So, we can utilize the some theory treating fingerprint spectra to analyze compound information in them.

In this paper, 24kind of pine samples belonging to Subgen.*Strobus* and Subgen.*Pinus* were quantitatively classified perfectly found on the two constants and the chemical compositions of their oleoresins. The results indicated that the new results both comply with that of classical classification, again exist obvious difference compared with that of classical classification. When using *P_g_* =61% as the theoretical standard, these 24 samples were clustered into one class/group,that is, Pinus. While when using *P_g_* = 70% as the theoretical standard, these 24 samples were divided into two Subgenus, that is, Subgen.*Strobus* and Subgen.*Pinus*. However, the results also showed that 3 samples out of 9 Subgen.*Strobus* (in terms of classically morphological classification), belong to Subgen.*Pinus* relying on the new method. Thus among these 24 samples, 18 of them belong to Subgen.*Pinus*, 6 of them were Subgen.*Strobus*. Moreover, by means of the characteristic sequences of these samples, the evolution relationship between the two subgenus could be revealed. The conclusion is Subgen.*Strobus* originated from Subgen.*Pinus*. On the other hand, the extremely large differences in geography and weather, can cause greater variation in chemical compositions of oleoresins of pines in the same Subgenus. This research uncovered that the two constants are qualified to be as the biological constants for Subgen.*Strobus* and Subgen.*Pinus*. Moreover, whether these two constants are suited for classifying other plants,and even used as category constants, such as species, genus, subgenus, family and so on.This is worth conducting much more researches.

## 2 Material

### 2.1 Oleoresin samples of Subgen.*Strobus* and Sugen.*Pinus*

The chemical composition characteristics come from oleoresin samples of 24 kinds of pines. These 24 oleoresin samples were listed in table 1.

**Table 1.**
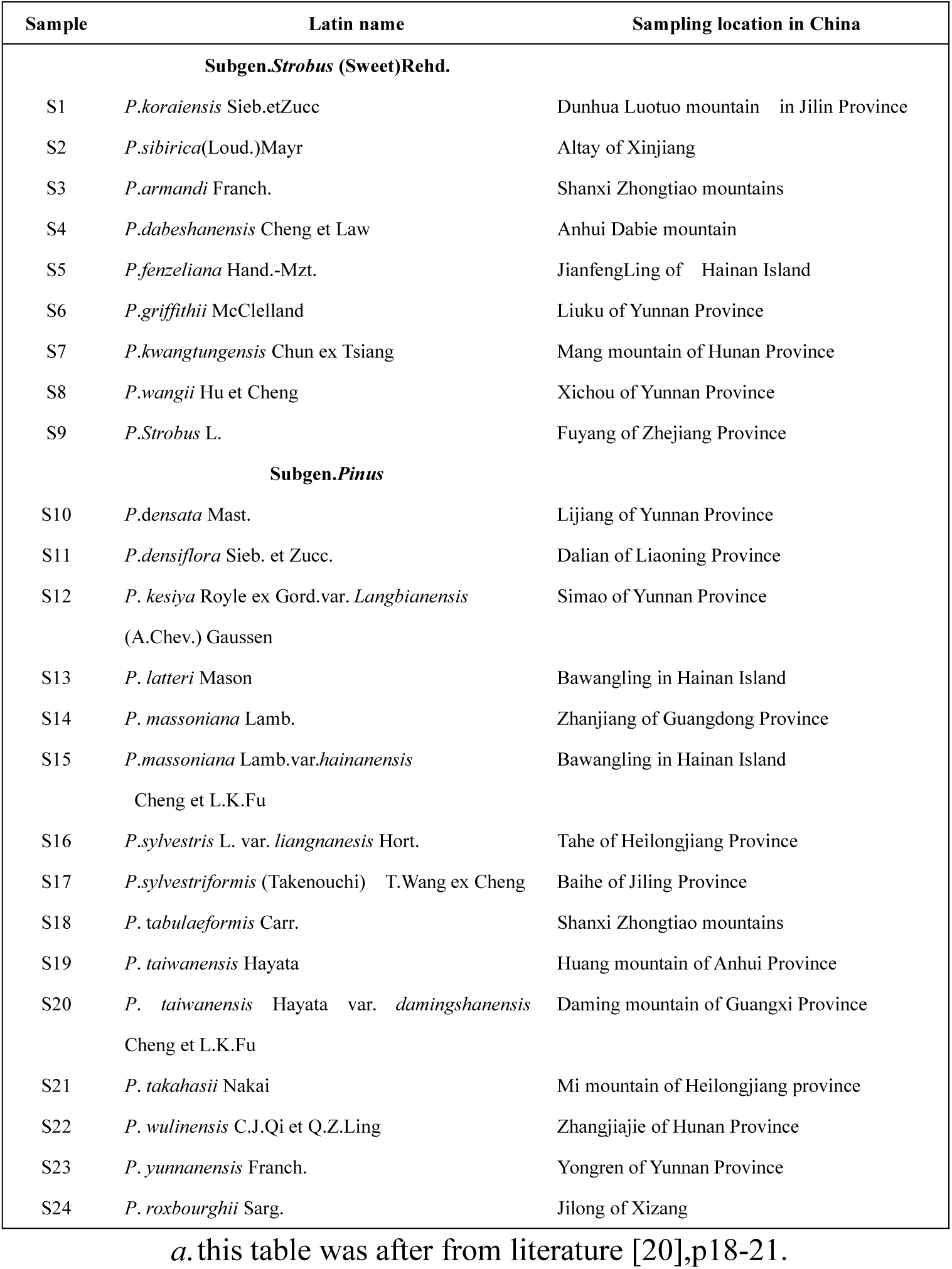
Oleoresin resources of 24 samples of Subgen.*Strobus* and Sugen.*Pinus*^a^

### 2.2 Compositions of oleoresins

From[20],the oleoresin compositions of 24 kinds of samples were obtained by means of methods in section 6, seen table 2-1,2-2,2-3 below.

**Table 2-1.** Compositions of oleoresins^a^

| Sample | The codes of compounds in oleoresins 1 to 19<sup>b</sup> |  |  |  |  |  |  |  |  |  |  |  |  |  |  |  |  |  |  |
| --- | --- | --- | --- | --- | --- | --- | --- | --- | --- | --- | --- | --- | --- | --- | --- | --- | --- | --- | --- |
|  | 1 | 2 | 3 | 4 | 5 | 6 | 7 | 8 | 9 | 10 | 11 | 12 | 13 | 14 | 15 | 16 | 17 | 18 | 19 |
| S1 | 1 | 2 |  | 4 | 5 |  | 7 | 8 | 9 | 10 | 11 |  | 13 | 14 | 15 | 16 | 17 | 18 | 19 |
| S2 | 1 | 2 |  | 4 | 5 |  | 7 | 8 | 9 | 10 | 11 |  |  |  |  |  |  | 18 | 19 |
| S3 | 1 | 2 |  | 4 | 5 | 6 |  | 8 |  | 10 |  | 12 | 13 |  | 15 | 16 | 17 | 18 | 19 |
| S4 | 1 | 2 |  | 4 | 5 |  |  | 8 |  | 10 | 11 |  | 13 | 14 | 15 | 16 | 17 | 18 | 19 |
| S5 | 1 | 2 |  | 4 | 5 |  |  | 8 |  | 10 |  |  | 13 | 14 |  |  | 17 | 18 | 19 |
| S6 | 1 | 2 |  | 4 | 5 | 6 | 7 | 8 |  | 10 | 11 |  |  | 14 |  |  |  |  | 19 |
| S7 | 1 |  |  | 4 | 5 |  | 7 | 8 |  |  | 11 |  |  |  |  |  |  |  | 19 |
| S8 | 1 | 2 |  | 4 | 5 |  |  | 8 |  | 10 | 11 |  |  |  |  |  | 17 | 18 | 19 |
| S9 | 1 | 2 | 3 | 4 | 5 | 6 |  | 8 |  | 10 |  | 12 |  |  |  |  |  |  | 19 |
| S10 | 1 | 2 | 3 | 4 | 5 |  | 7 | 8 | 9 | 10 |  |  | 13 |  |  | 16 | 17 | 18 | 19 |
| S11 | 1 | 2 |  | 4 | 5 |  |  | 8 | 9 |  |  |  |  |  |  | 16 | 17 | 18 | 19 |
| S12 | 1 | 2 |  | 4 | 5 |  |  | 8 |  | 10 |  |  |  |  |  | 16 | 17 | 18 | 19 |
| S13 | 1 | 2 |  | 4 | 5 |  | 7 | 8 |  | 10 |  |  |  |  |  |  | 17 |  |  |
| S14 | 1 | 2 |  | 4 | 5 |  | 7 | 8 |  | 10 |  |  |  |  |  | 16 | 17 | 18 | 19 |
| S15 | 1 | 2 | 3 | 4 | 5 |  | 7 | 8 |  | 10 |  |  | 13 |  |  | 16 | 17 | 18 | 19 |
| S16 | 1 | 2 | 3 | 4 | 5 |  | 7 | 8 | 9 | 10 |  |  | 13 |  |  |  |  | 18 |  |
| S17 | 1 | 2 |  | 4 | 5 |  |  | 8 |  | 10 |  |  |  |  |  | 16 | 17 | 18 | 19 |
| S18 | 1 | 2 | 3 | 4 | 5 | 6 |  | 8 |  | 10 |  |  | 13 |  |  | 16 |  | 18 | 19 |
| S19 | 1 | 2 |  | 4 | 5 | 6 |  | 8 |  | 10 |  |  |  |  |  | 16 | 17 | 18 | 19 |
| S20 | 1 | 2 |  | 4 | 5 | 6 |  | 8 |  | 10 |  |  |  |  |  | 16 | 17 | 18 | 19 |
| S21 | 1 | 2 |  | 4 | 5 | 6 |  | 8 | 9 | 10 |  |  |  |  |  | 16 | 17 | 18 | 19 |
| S22 | 1 | 2 |  | 4 | 5 | 6 |  | 8 | 9 | 10 |  |  |  |  |  | 16 | 17 |  |  |
| S23 | 1 | 2 |  | 4 | 5 |  |  | 8 |  | 10 |  |  | 13 |  |  | 16 | 17 | 18 | 19 |
| S24 | 1 | 2 |  | 4 | 5 |  |  | 8 |  |  |  | 12 | 13 | 14 |  |  |  | 18 |  |

**Table 2-2.**
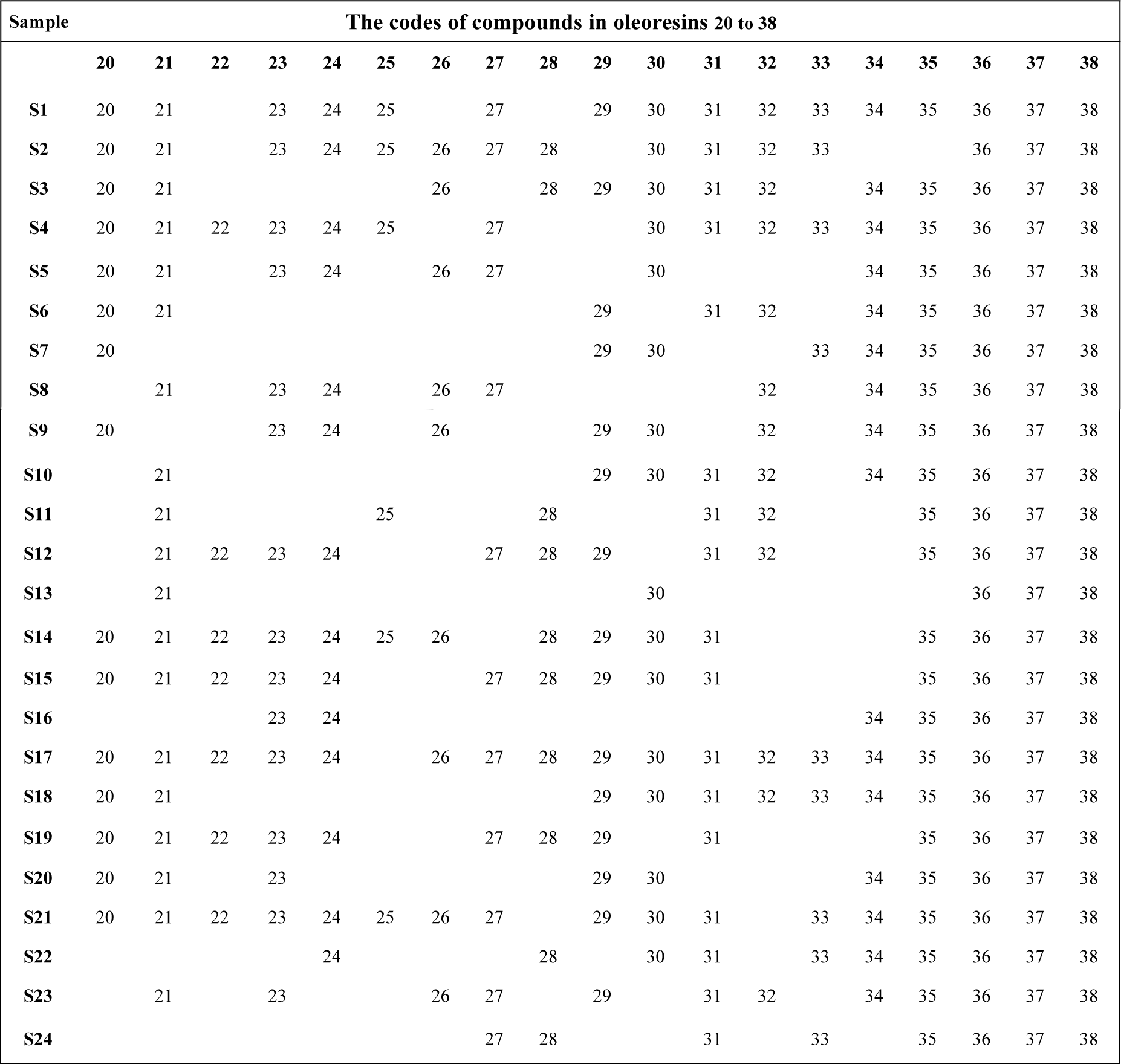
Compositions of oleoresins

**Table 2-3.**
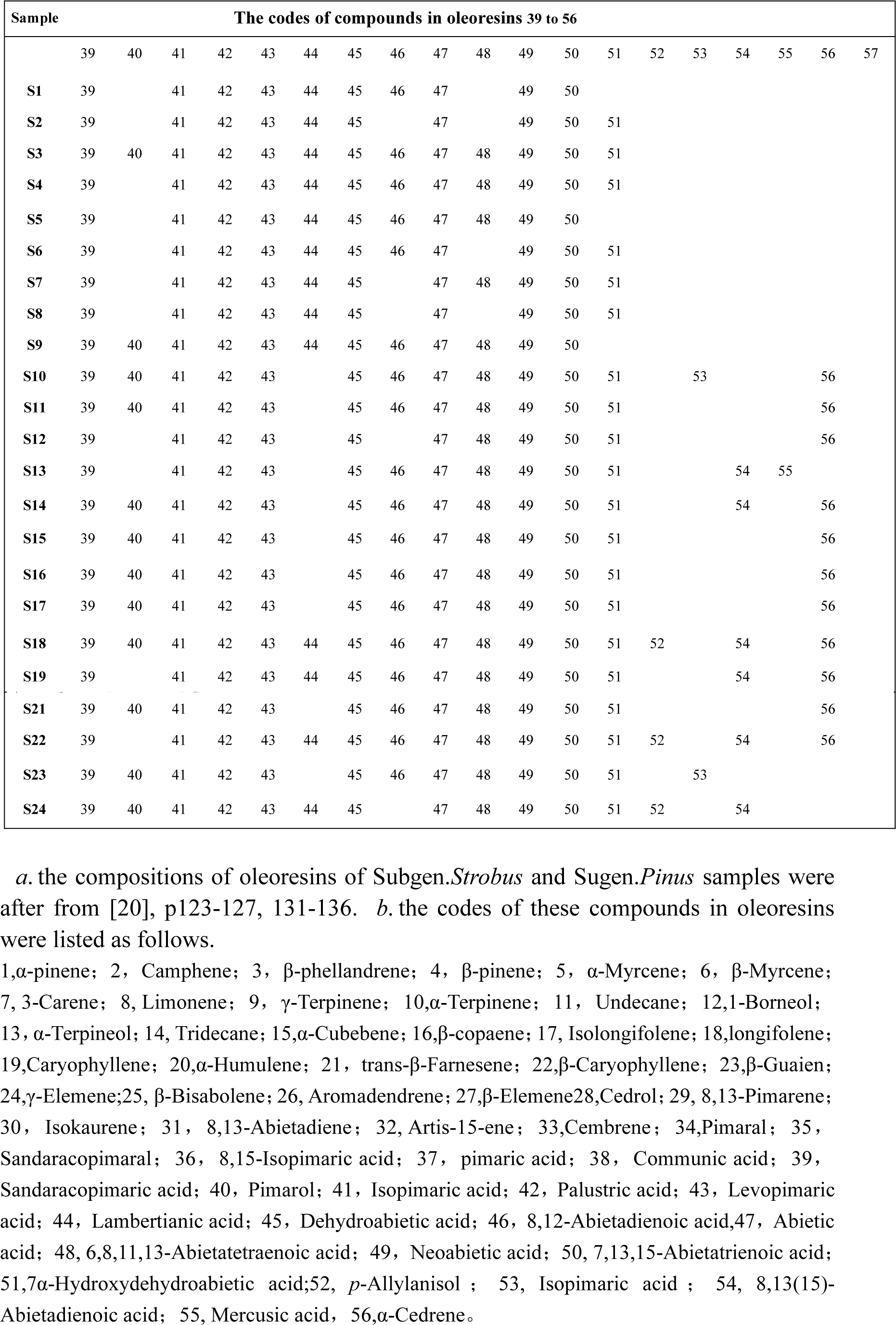
Compositions of oleoresins

## 3 Dual index sequence analysis of oleoresin samples

### 3.1 Constructing dual index sequence of different oleoresins

To apply dual index sequence analytical method [27–33], and to carry out the analysis based on oleoresin compositions. This method is briefly elucidated as follows.To calculate common composition ratios and variation ratios of the 24 oleoresins. Firstly, to select one sample being as the standard or reference, secondly, to calculate common composition ratios and variation ratios of all others to this reference. Then to arrange these samples in terms of the order that common composition ratio is from high to low value. For each sample by this way, one binary sequence with sample symbols, common composition ratios and variation ratios can be achieved. That is dual index sequences. The dual index sequences of 24 oleoresin samples were showed in **supplementary 1**.

### 3.2 Analysis on dual index sequences

#### 3.2.1 two constants from biological heredity and variation information theory

This biological common heredity and variation information theoretical equation[12], so-called dual index information theoretical equation, is displayed below.

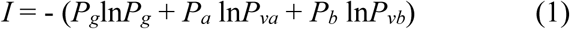

By means of this equation,we are able to calculate the mutual-action information value between any two organisms or any two evolution stages of a biological system.

The definitions of all variables and coefficients are showed there,according to literature[27–33].

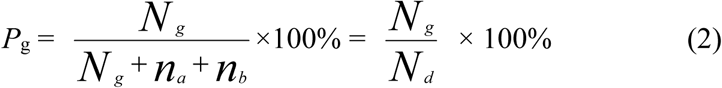

Common composition ratio *P_g_*:the ratio of common compositions *N_g_* existed in any two oleoresin samples *a,b* to the independent compositions *N_d_* in the *a*,*b*. The number of *N_d_* is equal to the kinds of different compounds in both sample *a*,*b*. *P*_g_ can be briefly expressed as P. This index P_g_ is the same as the Jaccard and Sneath, Sokal coefficients [9] intrinsically. *n_a_*, *n_b_* are the variation compositions in oleoresin sample *a*,*b*, respectively.

Other parameters are presented as follows.

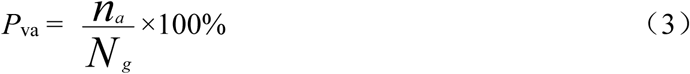

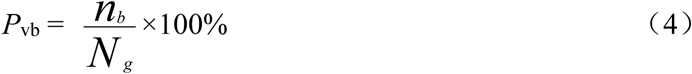

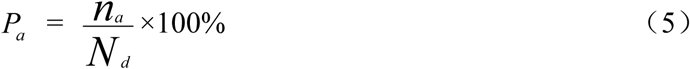

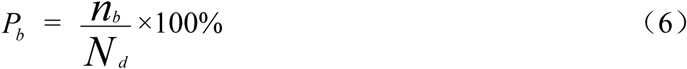

*P_va_* and *P_vb_* are the variation composition ratios of sample *a*,*b*, respectively.

*P_a_* and *P_b_* are the ratios of *n_a_* and *n_b_* to *N*_d_, respectively. They means the existed probabilities of *n_a_* and *n_b_*. More importantly, these parameters *P*_g_, *P_va_*, *P_vb_*, *P_a_* and *P_b_* are all established based on variables measured in experiments, without any coefficient determined by empirical knowledge. In fact, this information can be represented as a function *I_b_* = *f* (*n*_a_, *n*_b_, *N_g_*).

The relationship among these variables and parameters are elucidated below.

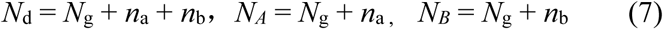

*N_A_* and *N_B_* are the number of compounds in sample *a*,*b*, respectively. The relationships of these variables may also be represented simply as follows.

**Fig. 1.**
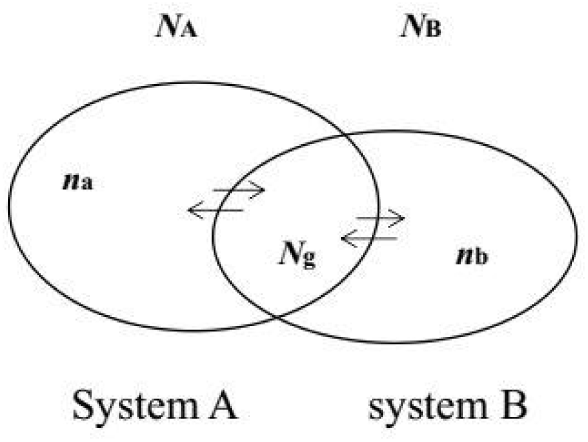
the relationships among different variables.

#### 3.2.2 To definite degree of symmetry

In order to research the symmetry and asymmetry of variation among any two biological systems,a new parameter was defined as *α*.

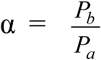, 0 ≦α ≦1. When α = 1, it shows two samples *a*,*b* are in symmetric variation state. When α =0, it expresses *a*,*b* are in absolute asymmetric variation state. When 0 ≦ α ≦ 1, it represents *a*,*b* are in different asymmetric variation states.

Depending on the two states, which are symmetric variation *n_a_ = n_b_*, *α =* 1, and asymmetric variation *n_a_* ≠ 0, *n_b_ =* 0, *α =* 0, with the maximum information values, two common composition ratios *P_g_*=0.610 and *P_g_*=0.695 can be achieved, respectively. Most interestingly, *P*_g_= 0.610 is very closed to gold ratio 0.618.

These two characteristic constants, corresponding to the two states *α =* 1, and *α =* 0, combing with the maximum number of effective sample method[13], are employed to analyze the 24 oleoresin samples based on their dual index sequences as in[12–15], the results were showed in section 3.2.2 and 3.2.3.

#### 3.2.3 Classification results relying on characteristic constant *P_g_* = 0.61 (61%)

The construction of characteristic sequences of samples is represented as follows.

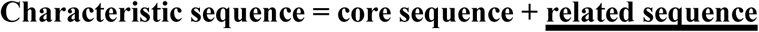

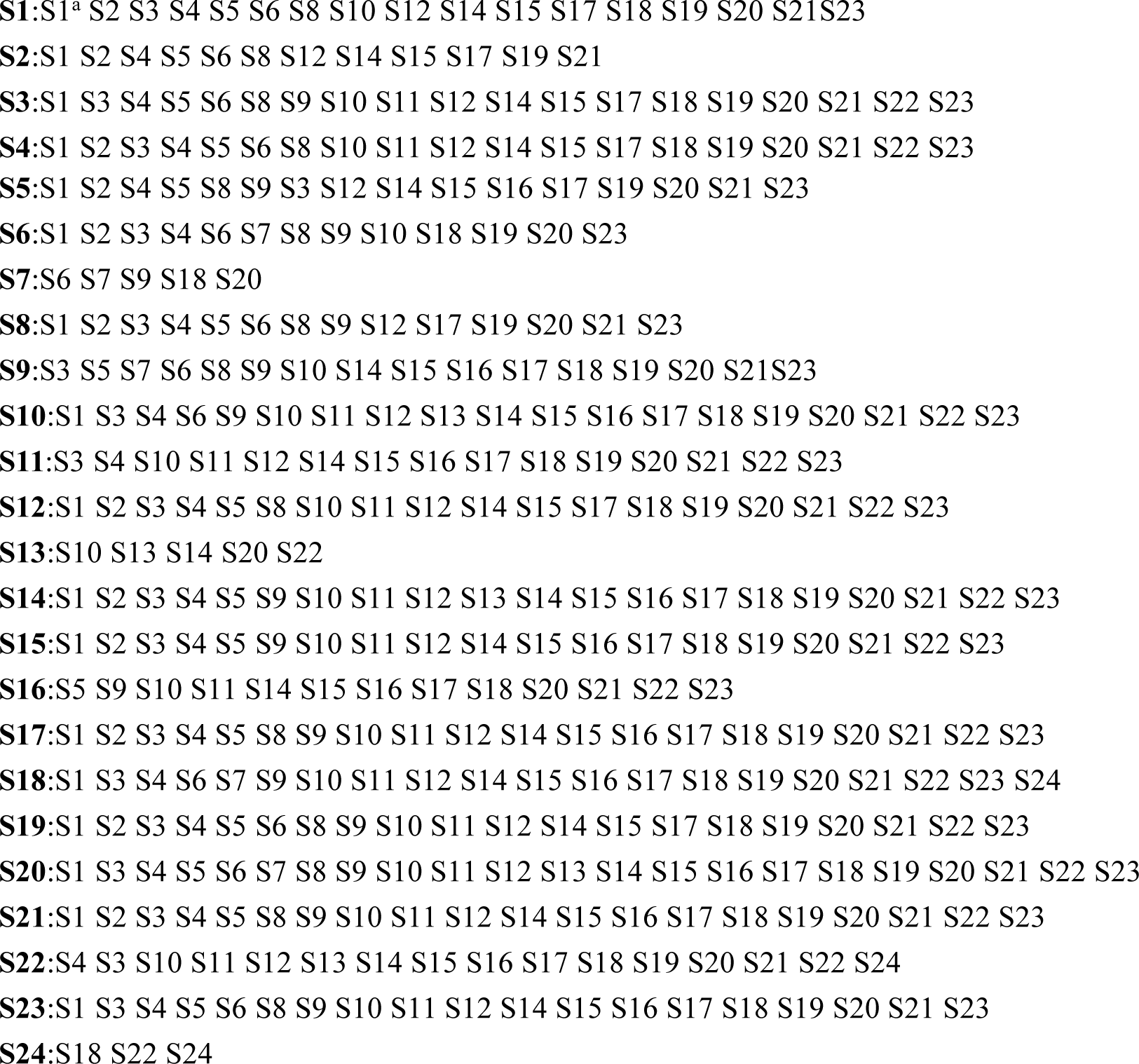

*a.* S1:S1^a^S2---,S1 belongs to its most similar sample, each sample belongs to its own most similar sample too.

Among these 24 samples, according to classically morphological classification results, S1~S9 are Subgen. *Strobus* pines, S10~S24 are Subgen. *Pinus* pines, which are all originated from China. The results based on *P*_g_ = 0.61= 61% were optimized. Samples in their characteristic sequences all belong to one group/set, there was no sample in related sequences. All samples in their core sequences were limited to one class, that is *Pinus*.

In accordance with characteristic sequence of each sample, S13 and S24 were of very short sequence, and differ from others greatly, although the 24 pines are all in one class. This may be because origin districts of S13 and S24 are unique. S13 is *P.latteri* Mason, it distributes in the district Hainan province of China, south of Guangdong province and south of Guangxi province in China, and indo-China penisula, the Malay penisula and the Philippines. These districts belong to tropical and subtropical areas, which are different from that of other pines obviously.

S24 belongs to Pinus roxburghii pine,one can know that it belongs to Subgen.*Pinus* by samples in its characteristic sequence. However, there were distinct differences compared to the characteristic sequences of other samples. This may result from that its distribution area is of great differences related to that of other samples. S24 originated from Jilong district in south of Tibet Plateau, where is at an altitude of 2 100 to 2 200 meters. The distribution center is at the southern slop of Himalaya mountain. On the other hand, it also originated from the kingdom of Bhutan, the Sikkim district, India, Nipper, even from Afghanistan. S24 come from Jilong, which is of significant differences compared to origin districts of other samples. This may leads to vast difference in compositions compared with other samples. So does the S13.

The results showed ahead expressed that *P_g_* ≧ (61±3)% could be employed as the absolute theory standard [14,15]to determine pines of Pinus in practice. This verified that common heredity and variation information theory equation fit to describe biological evolution rules.

#### 3.2.4 Classification results based on characteristic constant *P*_g_ =0.70

In order to distinguish Subgen. *Strobus* from Subgen. *Pinus*, *P_g_* =70% was accepted as the absolute theory standard, and practical criterion *P_g_* ≧ (70±3)% [14,15] was used to be classification standard, combing the maximum number of effective sample method[13]. By practical analysis, the optimized results were obtained at common composition ratio *P*_g_ = 69%,and these results were listed in table 3.

**Table 3.**
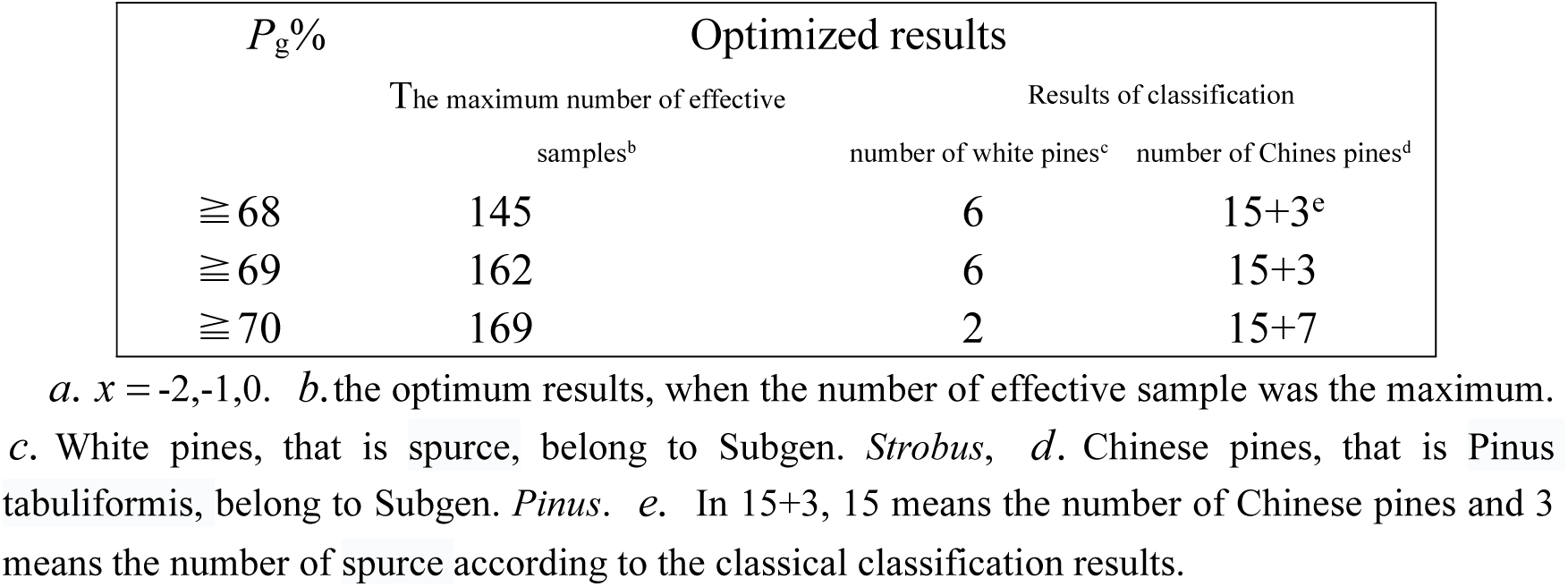
The number of effective samples based on P_g_≧(70 ± x a)%

According to table 3, although the number of effective samples was the maximun, when *P_g_* ≧ 70%, the results were not correct, since there were only 2 samples belonging to Subgen. *Strobus*, 22 samples were Subgen. *Pinus.* This case is overclassification.

Combining with the maximum number of effective sample method[13],

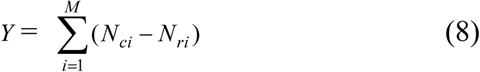

*Y* : the maximum number of effective samples in characteristic sequences of sample set.

*N_ci_* : the number of samples in the *i* th samples’ characteristic sequence.

*N_ri_* : the number of samples in *i* th samples’ related sequence.

*M* : the number of total samples.

*Y* reflects degree of effective classification of sample set. The larger the *Y,* the more clear the classification, and the much less the samples in related sequences. This means that the more samples in core sequences, the classification is more ideal.The number of the most ideally effective samples is 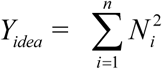, in which *N_i_* is the number of samples in *i* th class, *n* is the number of classes. In this case, the *N_ci_* is equal to that of samples in a class,and *N_ri_* =0, tending to ideal classification. For instance, in this study, the number of samples belonging to Subgen*.Strobus* is 6, and that of samples belonging to Subgen.*Pinus* is 18, then the *Y_idea_* = 6^2^ + 18^2^ = 360. This method is an excellent one for quantitatively judging whether the result is pros and cons. At the same time, a big advantage for this approach is that it can avoid over classification, compared to other pattern recognition methods,and assure the result is objective classification compared with other methods. On the other hand, in this research the *Y* is obviously lower than *Y_idea_*, it indicated there are great variation among samples. Depending on the compounds in oleoresins, classification results of these 24 samples of Subgen*.Strobus* and Subgen.*Pinus* were showed as follows.

The construction of characteristic sequences of the 24 samples is :

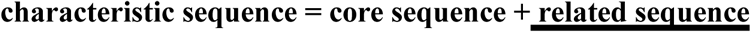

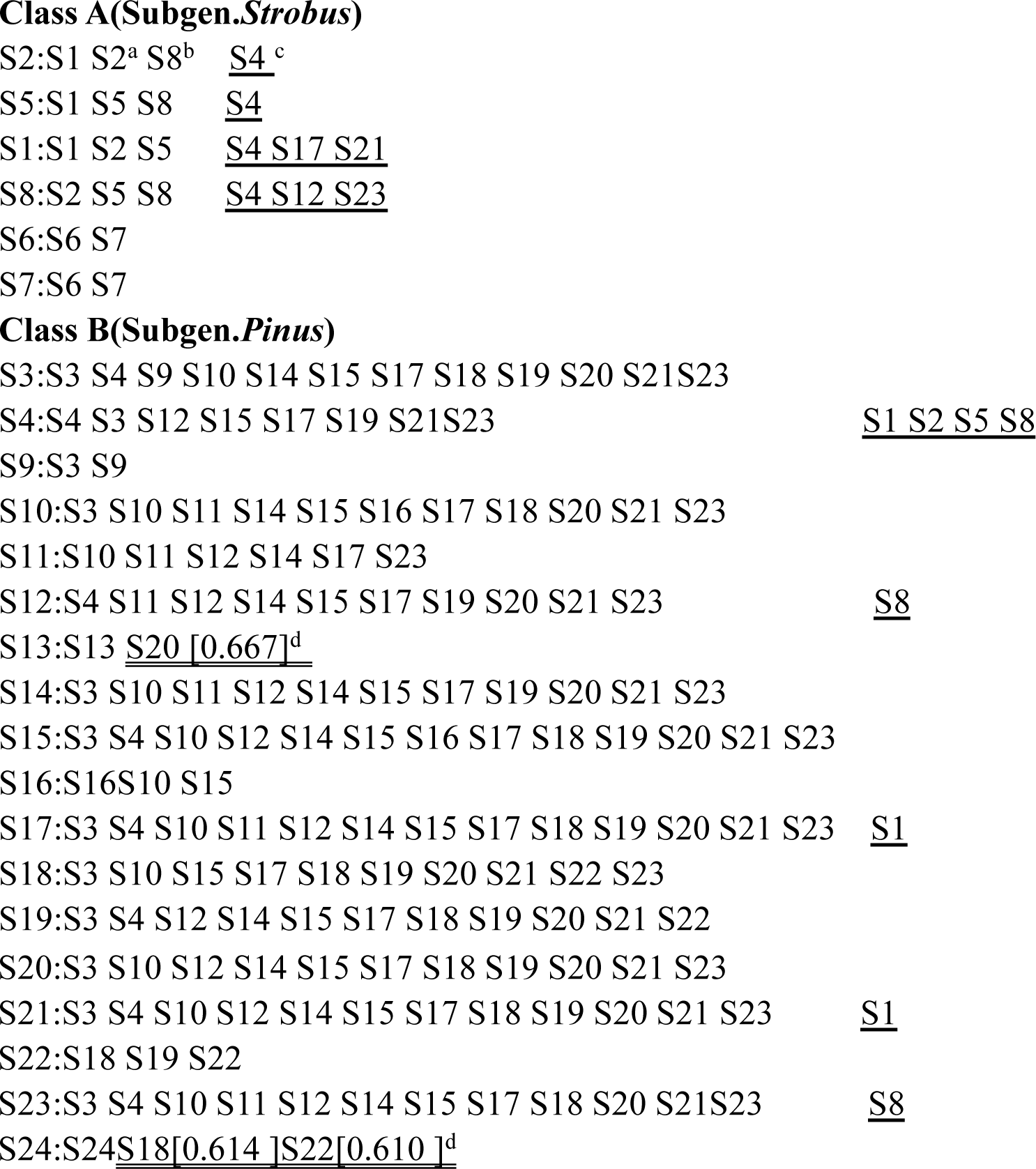

S2:S1 S2^a^S8^b^S4^c^, this sequence was the characteristic sequence of S2. a. S2, each sample is the most similar sample in its own characteristic sequence. b. S1S2S8 was the core sequence of S2. c. S4 was the sample in related sequence of S2. d. S13:S13S20[0.667], 0.667 was the common composition ratio between S13 and S20. S24:S24S18[0.614]S22[0.610], the part sequence with double underline was the close sequence of S24, in which the composition ratios of sample S18,S22 related to S24 were lower than 69%. This part sequence shows that S24 belongs to Subgen.*Pinus*.

## 4 Evolution pattern analysis of the two subgenus

In above characteristic sequences. the common composition ratios of samples to reference samples were equal to or larger than 69%. These results represented that Subgen.*Strobus* and Subgen.*Pinus* could be distinguished clearly with the standard P_g_ ≧ 69%. In class A Subgen.*Strobus*, the characteristic sequences of S1,S2, S5, S8 were with obviously related sequence made up of samples of Subgen.*Pinus*. This uncovered that Subgen.*Strobus* is greatly of the properties of Subgen.*Pinus*. The characteristic sequences of samples of Subgen.*Strobus* included samples of Subgen.*Pinus*, and the ratios of Subgen.*Strobus*’ samples to Subgen.*Pinus*’ samples were very high, and variations among characteristic sequences of Subgen.*Strobus* were distinct. Compared with Subgen.*Pinus*’ samples, their characteristic sequences almost contain no Subgen.*Strobus*’ samples. This asymmetry revealed that Subgen.*Strobus* should evolved from Subgen.*Pinus*. These relationships were displayed qualitatively in figure 2.

**Fig. 2.**
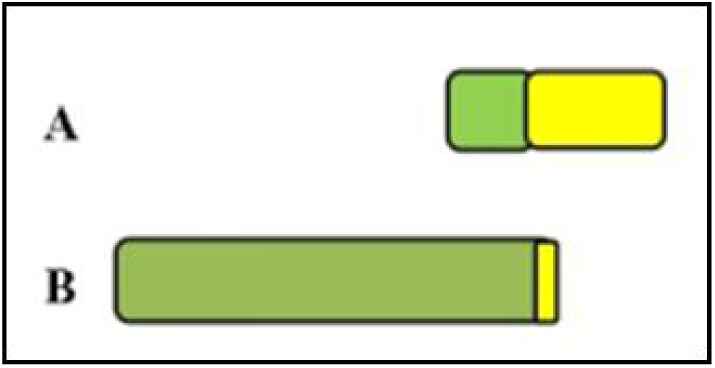
the sample set of characteristic sequences of Subgen.*Strobus* and Subgen.*Pinus*. A. Subgen.*Strobus*. B. Subgen.*Pinus* Green/deep color region express Subgen.*Pinus*. yellow/light color region express Subgen.*Strobus*. The large size of area show the number of samples was large.

On the other hand. in characteristic sequences of Subgen.*Strobus*,the number of samples in core sequence of every Subgen.*Strobus* sample was small. and there were great change among their core sequences. This fact pointed out there were obvious variations among them. and the origins should be from multiple sources.

From geographic space point of view, if Subgen.*Pinus* is in a concentration distribution space, Subgen.*Strobus* should distribute around the space of Subgen.*Pinus*. In the evolution chain,Subgen.*Pinus* was first originated from it ancestor, then Subgen.*Strobus* came from Subgen.*Pinus*. This involving relation existed in characteristic sequences was the equal of the overlapping relation between different sets. By this view, the origins of different organisms could be inferred from the mutual involving relation, and evolution history could be portrayed briefly. This verified that the common heredity and variation information theoretical equation could discover the quantitative evolution principle of biology, and it provides a new theory for investigating biological evolution.

From the related sequences of Subgen.*Strobus*’ samples,the samples in the related sequences were mainly S4,S12,S17, S21 and S23, which are Subgen.*Pinus*. This inferred that Subgen.*Strobus* were mainly evolved from S4,S12,S17, S21 and S23 of Subgen.*Pinus*. Grounded on the core sequences of Subgen.*Strobus*’ samples, they can be divided into two subgroups, subgruop1:S1,S2,S5,S8 and subgroup 2: S6,S7. Their core sequences construct two independent sets. This showed there are obvious differences in property between the two subsets.

S4, S12, S17, S21 and S23 existed in related sequences of S1,S2,S5,S8, this manifested there exist the kinship between S1,S2,S5,S8 and Subgen.*Pinus*.

In accordance with characteristic sequences, samples S3,S4,S9, which are Subgen.*Strobus* based on the classical classification results, could be classified into Subgen.*Pinus* perfectly by means of this new approach.

On the other flip side, there were no or few samples in the related sequences of S3,S4,S9~S24.This revealed that there are distinct differences between Subgen.*Strobus* and Subgen.*Pinus*. Depending on the involving relation between characteristic sequences of Subgen.*Strobus* and subgen. Pinus. The maximum likelihood is Subgen.*Strobus* evolving from Subgen.*Pinus*. This also means Subgen.*Pinus* is ancestor, Subgen.*Strobus* is the son.

The results offered above represented that P_g_≧ 69% could be used as the absolutely quantitative standard for distinguishing Subgen.*Strobus* and Subgen.*Pinus*. Two subgenus are of significant asymmetry variation in evolution. No mater from the common composition ratio values, or from the properties, the conclusion are all fit to theoretical derivation. This demonstrates that dual index information theory equation is capable of accurately describing biological evolution rules.

In Subgen.*Pinus*, some kinds of pine species were of distinct feature. Although S4 was divided into class B, that is Subgen.*Pinus*, its related sequence contained many samples of Subgen.*Strobus*, S4:S3S4S12S15S17S19S21S23S1S2S5S8, this expressed S4 is in the transition stage of evolution from Subgen.*Strobus* to Subgen.*Strobus*. S4 may be named the transition species from Subgen.*Pinus* to Subgen.*Strobus*.

Although S9 was Subgen.*Pinus*, there were great differences between S9 and other samples of Subgen.*Pinus*, based on its characteristic sequence. S9 is Pinus strobus (eastern white pine), the geographic feature of its origin is extremely different from that of other samples. This brings about its variation being greatly compared to other samples. Pinus strobus Linn originated from eastern north of America, its most similar sample was S3, that is Pinus armandii Franch, which originate from Hua mountain in Shanxi province of China. The geographic feature of origin districts is similar with each other. The climate is moist and warm cool.

S13 is Pinus *Latteri* Mason, seen in section 3.2.2. Its geographic areas differ from that of other pine species. This resulted in its properties, chemical compositions being of unique feature. From its characteristic sequence, its most similar sample was S20, whose common composition ratio was merely equal to 67%, less than 69%. Thus S13 could be viewed as an independent class.

S20 is pinus taiwanensis Hayata var, it distributes in Daming mountain of Guangxi province and Fanjing mountain of Guizhou province in China. Its geographic features differ from that of other pinus origins significantly.

S24 is Pinus roxburghii (Chir Pine, or longleaf lndian) pine, and belongs to Subgen.*Pinus,*depending on samples in characteristic sequence. However, there existed obvious differences between it and other pine species of Subgen*.Pinus.* This was because of the extreme difference in its growth geographic environment between it and other samples of Subgen.*Pinus*. Its origin center is in the southern slop of Himalaya mountains. S24 was collected from this center,its unique geographic environment makes it differ from other samples of Subgen.*Pinus* significantly, and determines that there was no sample with high similarity to it. The most similar samples to it were S18, S22, while the common composition ratios are merely 0.614 and 0.610, which are far less than 69%, respectively.

Compared the characteristic sequence of classes A with that of class B, one can find out that most characteristic sequences of samples in Subgen.*Pinus* vary slightly. This indicated that Subgen.*Pinus* is in steady evolution state. In contrast, the characteristic sequences of Subgen*.Strobus* samples change greatly and randomly.These revealed that Subgen*.Strobus* is in unsteady evolution state. The analytical results expressed that the characteristic constant *P_g_* =70%, obtained from dual index information theory equation, relying on its asymmetric variations, combing with the maximum number of effective sample method[13], could be employed as absolutely theoretical standard for classifying Subgen*.Strobus* and Subgen.*Pinus*. Another constant *P_g_* =0.61, based on symmetric variation, could be as the absolute theoretical standard for determining which trees are Pinus.

## 5 Conclusion

The two constants derived from biological common heredity and variation information theory equation, combining with compounds in oleoresins metabolized by pines, could accurately identify Subgen.*Strobus* and Subgen.*Pinus* without any help of empirical knowledge related to learning samples. This showed that common heredity and variation information theory equation can uncover biological evolution rule excellently. The two constants can reveal the features of some biological categories. In other hand, the characteristic sequences of samples in each class were able to express evolution information of organisms. The involving relationships between core and related sequences, which belong to different classes, respectively, can uncover evolution history and relationship of different classes. One can determine which is ancestor, and which is the son. This research indicated that Subgen.*Strobus* should originate from Subgen.*Pinus* and come from multiple species of Subgen.*Pinus*. Common heredity and variation information theory provides us with a new approach for accurately classifying biology, analyzing evolution relationships of biology by applying molecular structure, molecular kind information. The two constants may be used as absolutely theoretical criteria for classifying different biological categories, and may revise the results obtained based on classical classification methods.For these categories, such as species, subgenus and genus with very close kinship, their inner similarity is high, the two constants can be adopted as the theoretical criteria for determining them. However, for many other categories, such as family,order, class,phylum, and kingdom, being of distant relatives, samples in one category of them are of low similarity. These two constants should not suit to these categories. For family, the two constants may fit to, or not match with the classification of them. Further more, whether biological common heredity and variation information theoretical equation is suitable for analyzing structure information in DNA, protein, all these need to be further investigated for us.

## 6 Instruments and methods[20]

To see supplementary 2.

